## Supplementary Materials for "Non-linear Archetypal Analysis of Single-cell RNA-seq Data by Deep Autoencoders"

### Supplementary Figures

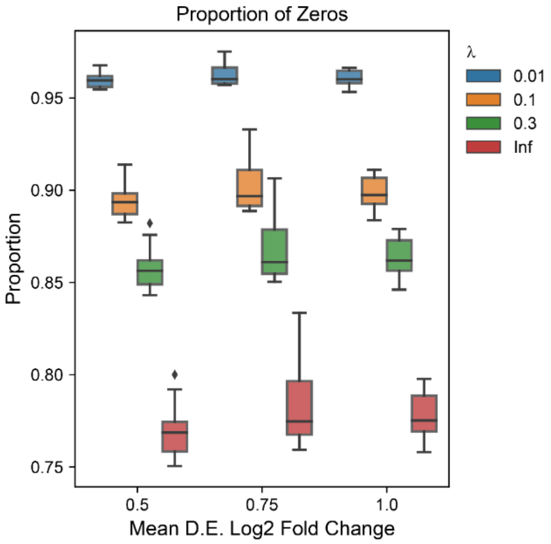


**Supplementary Fig. 1** Dropout proportions of synthetic scRNA-seq datasets across different values of $\lambda$ and mean differential expression (D.E.) log2 fold change of GEP-specific genes. Each box and whisker plot was plotted based on ten simulated datasets. Central lines represent medians, boxes represent the interquartile range (IQR), and the upper/lower whisker represents the largest/smallest value no further than 1.5 × IQR.

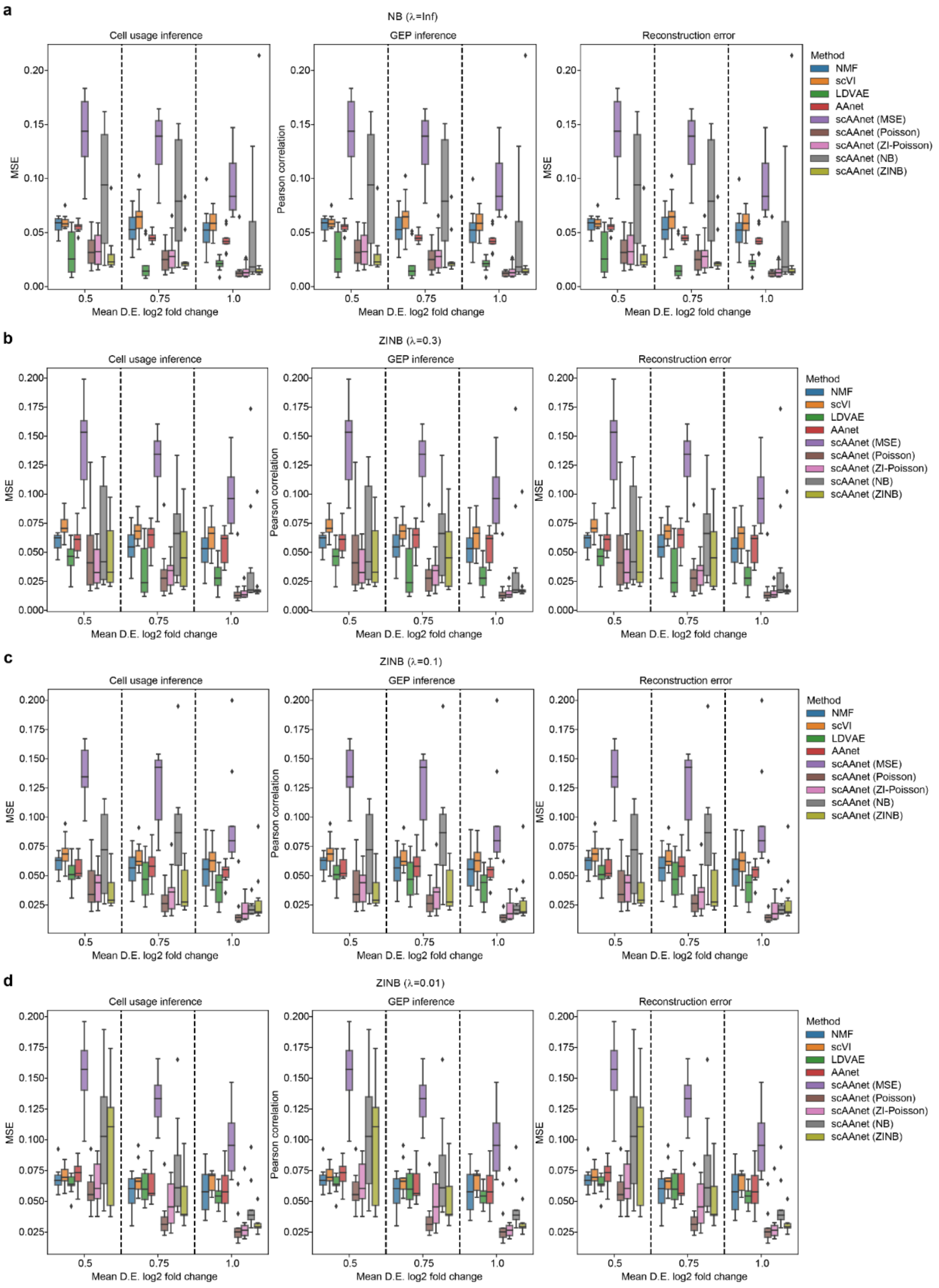


**Supplementary Fig. 2** Performance of scAAnet and other methods on synthetic scRNA-seq datasets that were simulated under NB (**a**) and ZINB distribution with $\lambda$ = 0.3 (**b**), $\lambda$ = 0.1 (**c**) and $\lambda$ = 0.01 (**d**). For each panel, figures from left to right are MSE between inferred cell usages and true usages of the 4 GEPs, Pearson correlation between inferred GEPs and true GEPs, and MSE between reconstructed scaled means and true means. Each box and whisker plot was plotted based on ten simulated datasets. Central lines represent medians, boxes represent the IQR, and the upper/lower whisker represents the largest/smallest value no further than 1.5 × IQR. D.E.: differential expression.

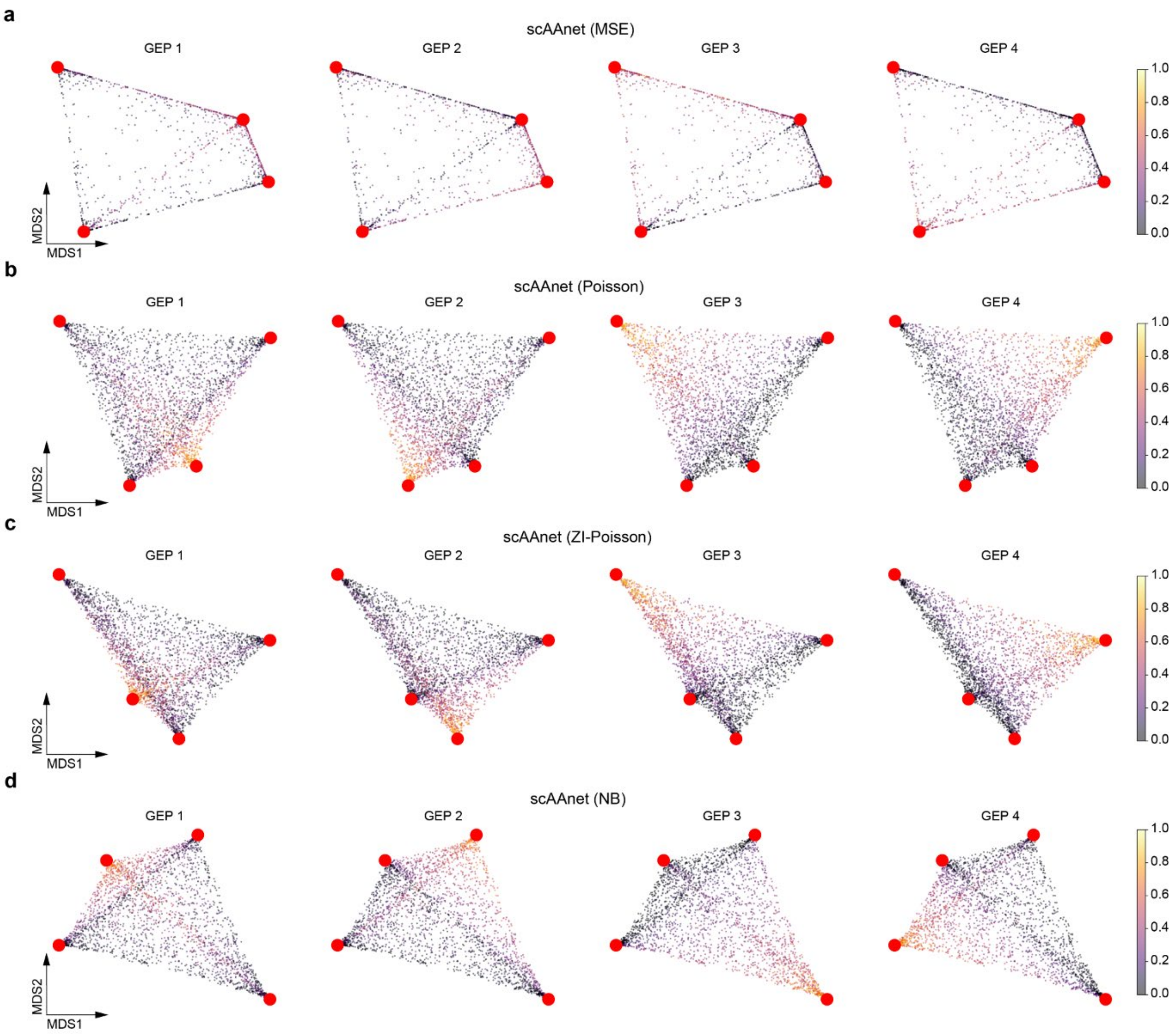


**Supplementary Fig. 3** MDS interpolation visualization of archetypal space recovered from scAAnet with MSE (**a**), Poisson (**b**), ZIP (**c**), and NB (**d**) distribution as the reconstruction error term. Red dots are the locations of archetypes. Figures in each column are colored by the true cell usage of the corresponding archetype (GEP).

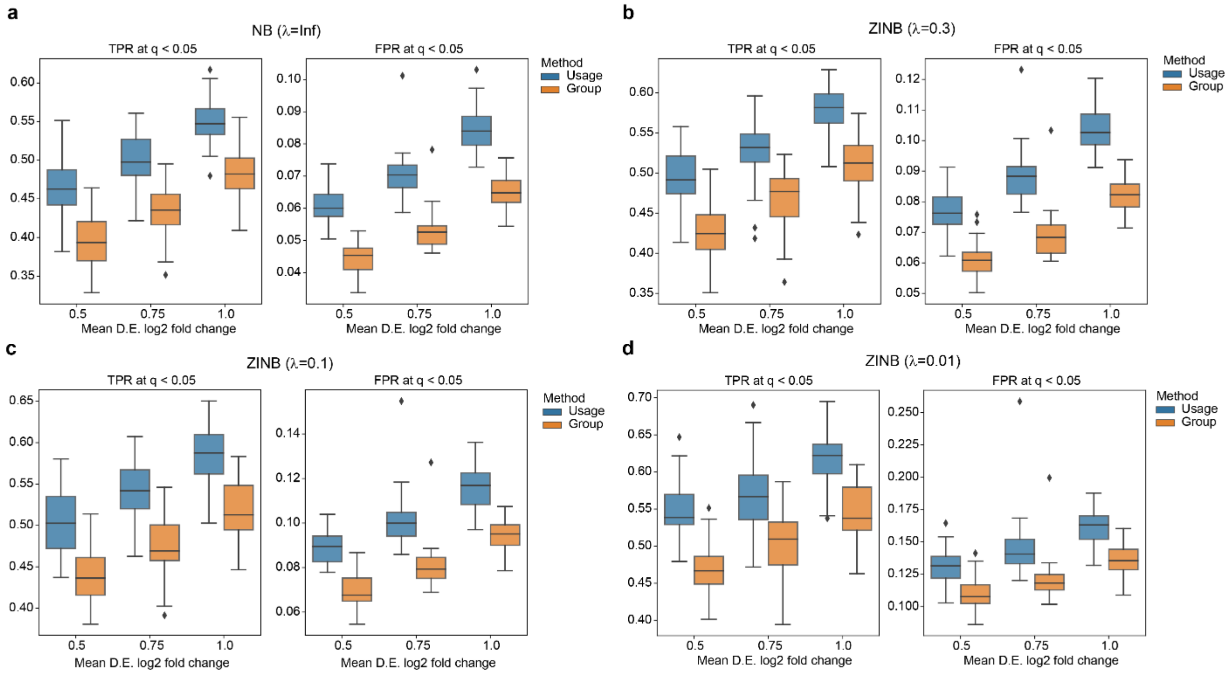


**Supplementary Fig. 4** Comparison of DEG test results between using continuous usages and discreate group assignment on synthetic scRNA-seq dataset that were simulated under NB (**a**) and ZINB distribution with $\lambda$ = 0.3 (**b**), $\lambda$ = 0.1 (**c**) and $\lambda$ = 0.01 (**d**). For each panel, figures from left to right are TPR at q-value < 0.05 and FPR at q-value < 0.05 calculated across different signal-to-noise ratio levels. Bonferroni correction was used to obtain q-values. Each box and whisker plot was plotted based on ten simulated datasets. Central lines represent medians, boxes represent the IQR, and the upper/lower whisker represents the largest/smallest value no further than 1.5 × IQR. TPR: true positive rate; FPR: false positive rate.

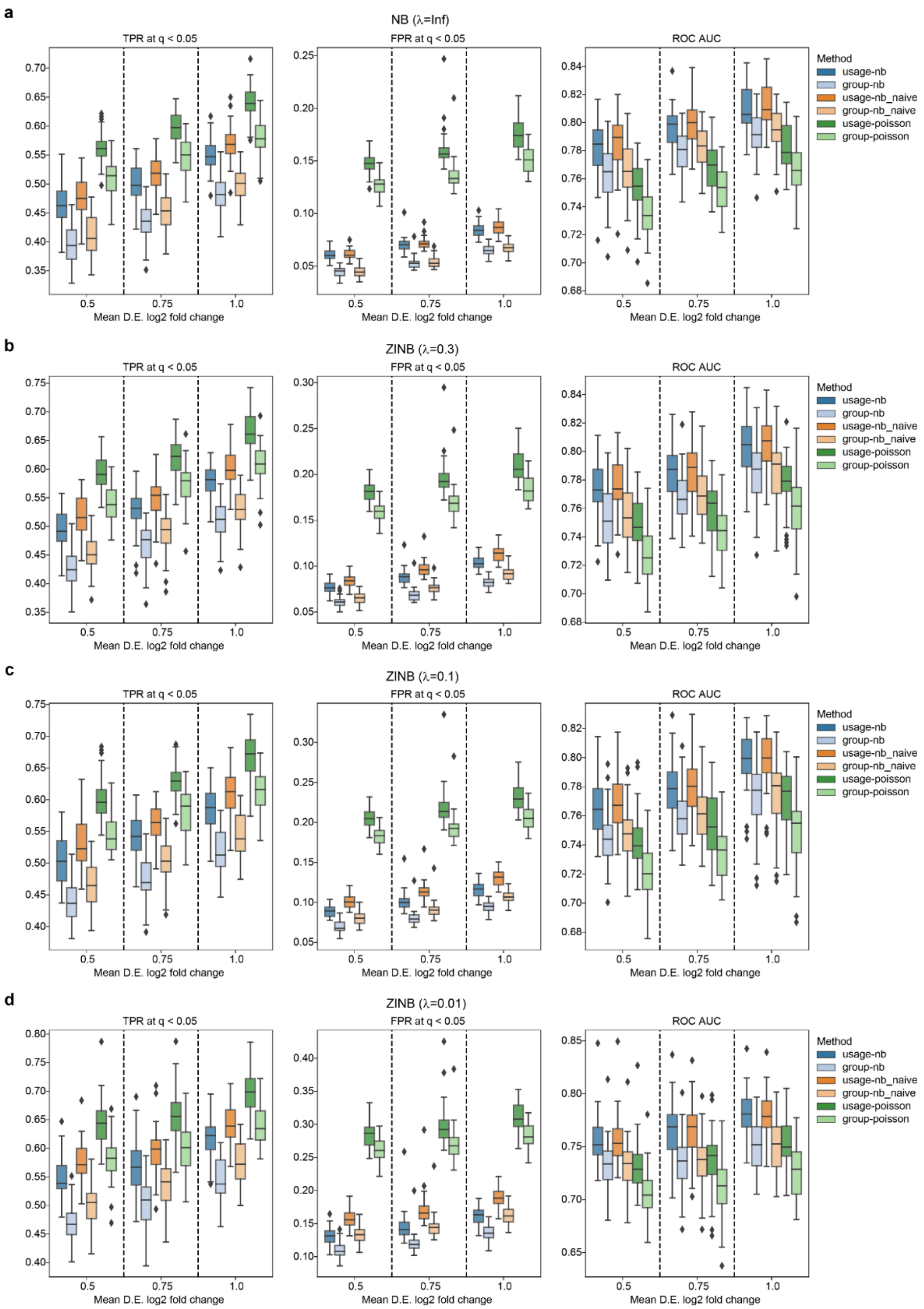


**Supplementary Fig. 5** Comparison of DEG test results among different regression models on synthetic scRNA-seq dataset that were simulated under NB (**a**) and ZINB distribution with $\lambda$ = 0.3 (**b**), $\lambda$ = 0.1 (**c**) and $\lambda$ = 0.01 (**d**). For each panel, figures from left to right are TPR at q-value < 0.05 and FPR at q-value < 0.05 calculated across different signal-to-noise ratio levels. Bonferroni correction was used to obtain q-values. Each box and whisker plot was plotted based on ten simulated datasets. Central lines represent medians, boxes represent the IQR, and the upper/lower whisker represents the largest/smallest value no further than 1.5 × IQR.

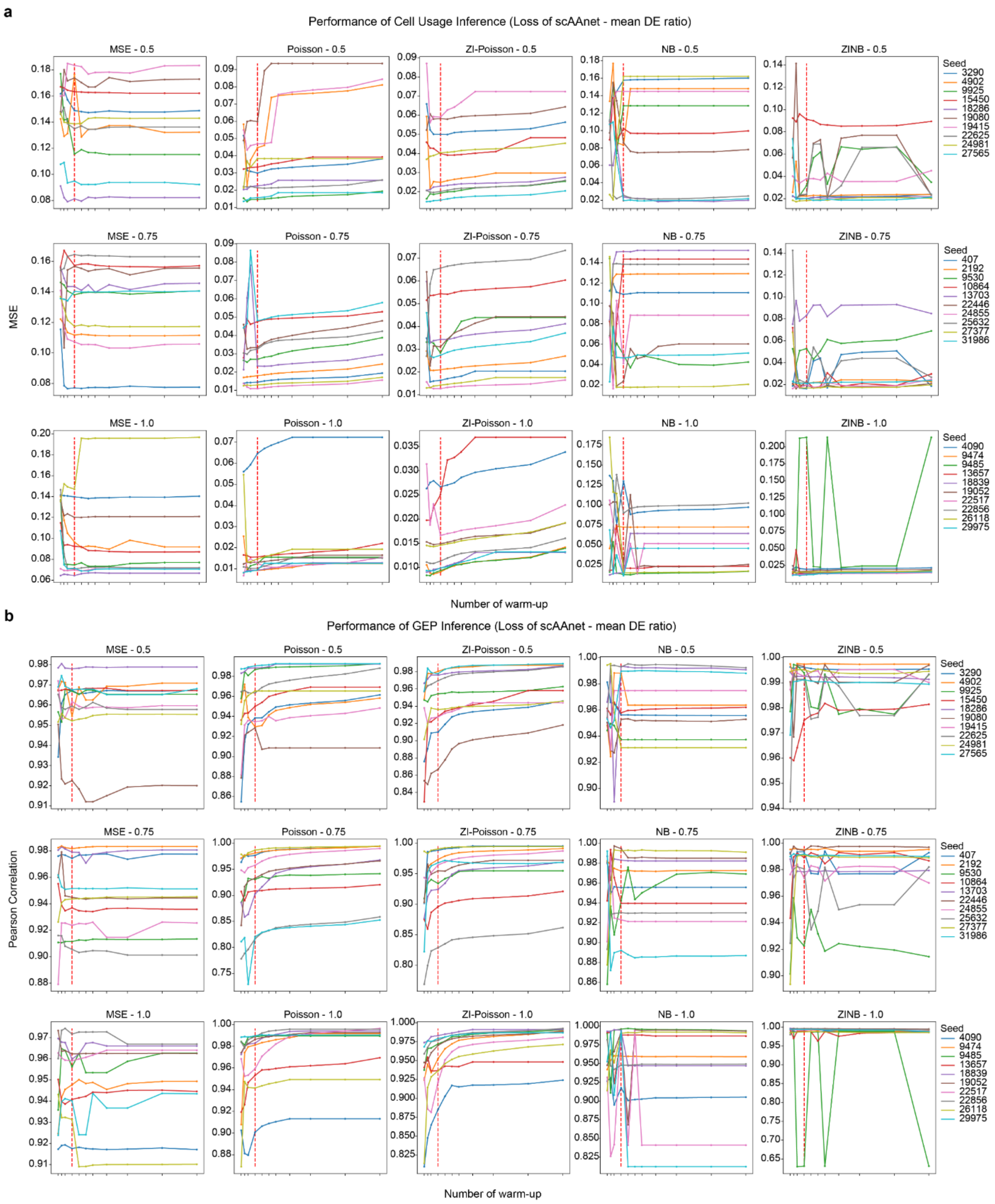


**Supplementary Fig. 6** Performance of cell usage inference (**a**) and GEP inference (**b**) over increasing number of warm-up.

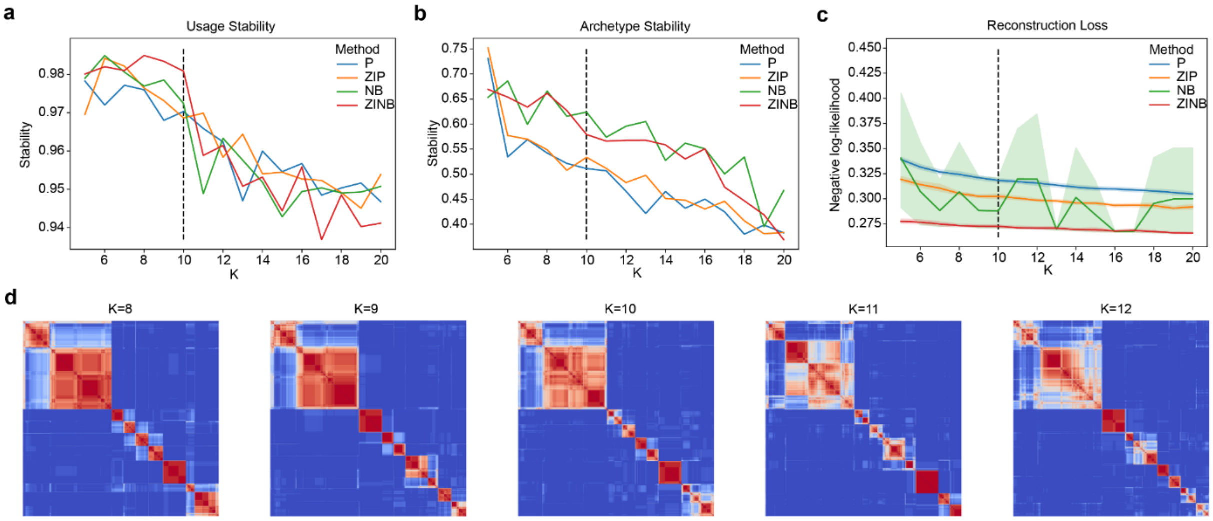


**Supplementary Fig. 7** Selection of K in the pancreatic islet dataset. **a**, **b**, **c** Stability of usage (**a**), stability of archetypes (**b**) and reconstruction loss (**c**) across different Ks for four count distributions. **d** Consensus clustering matrices showing the usage stability under K = 8-12.

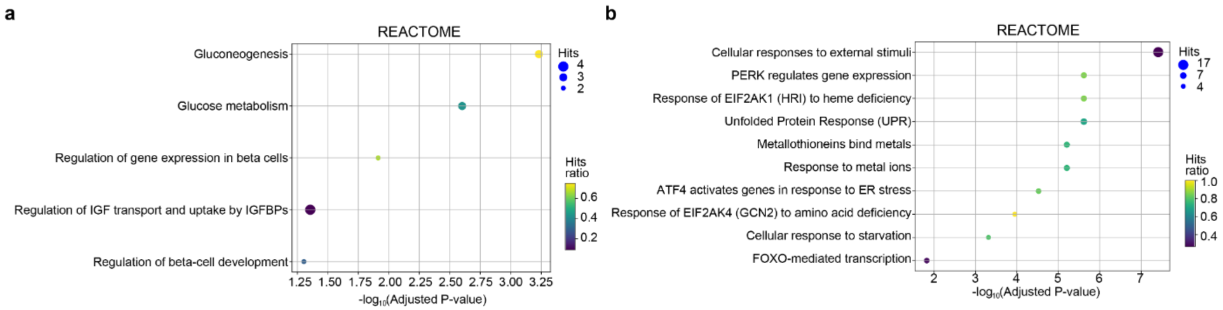


**Supplementary Fig. 8** REACTOME enrichment results of the two GEPs related to beta cells in the pancreatic islet dataset. **a** Top enriched terms identified using significantly upregulated genes in GEP 2. **b** Top 10 enriched terms identified using significantly upregulated genes in GEP 7.

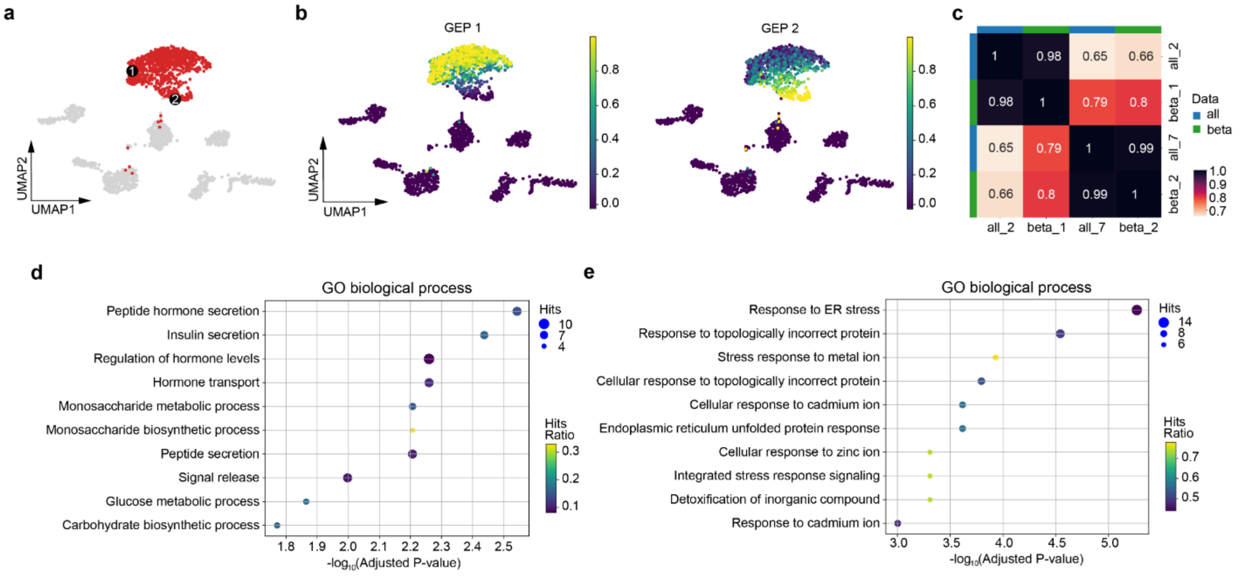


**Supplementary Fig. 9** scAAnet is able to identify two GEPs using beta cells only that are similar to GEP 2 and GEP 7 identified in the pancreatic islet dataset. **a** The UMAP visualization of the pancreatic islet scRNA-seq dataset, with beta cells highlighted in red. Black dots are locations of cells that have the largest usage of the 2 GEPs (marked in Arabic numerals). **b** UMAPs colored by inferred cell usage for each GEP. **c** Heatmap showing the Pearson correlations among GEP 1 and GEP 2 identified using beta cells only and GEP 2 and GEP 7 identified using the entire dataset. **d** The top 10 enriched GO biological process terms using 35 significantly upregulated genes of GEP 1. **e** The top 10 enriched GO biological process terms using 130 significantly upregulated genes of GEP 2.

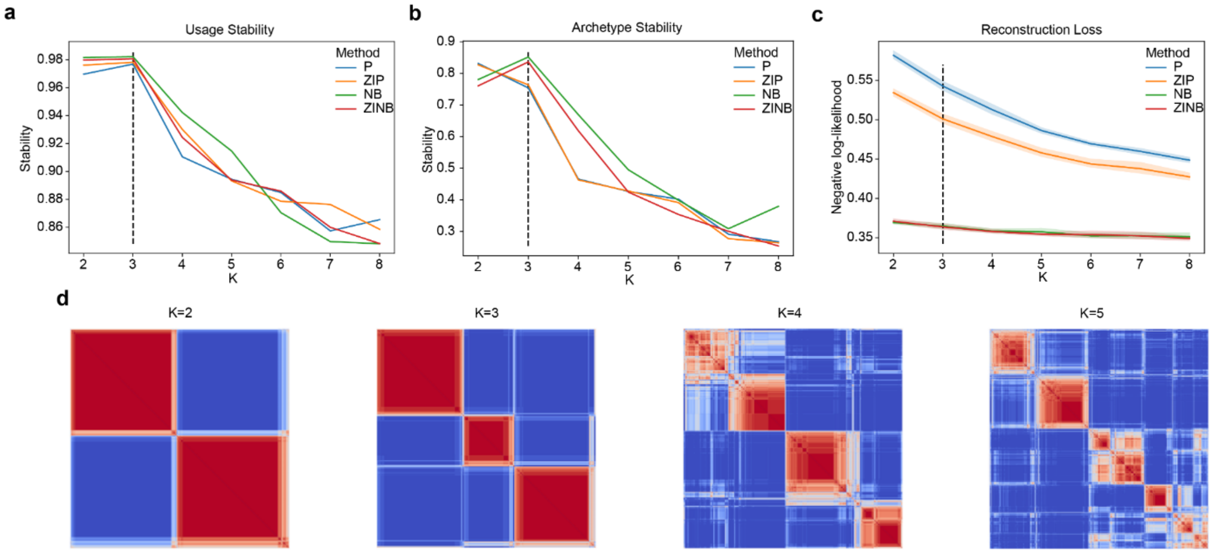


**Supplementary Fig. 10** Selection of K in the lung fibroblast and myofibroblast cells. **a**, **b**, **c** Stability of usage (**a**), stability of archetypes (**b**) and reconstruction loss (**c**) across different Ks for four count distributions. **d** Consensus clustering matrices showing the usage stability under K = 2-5.

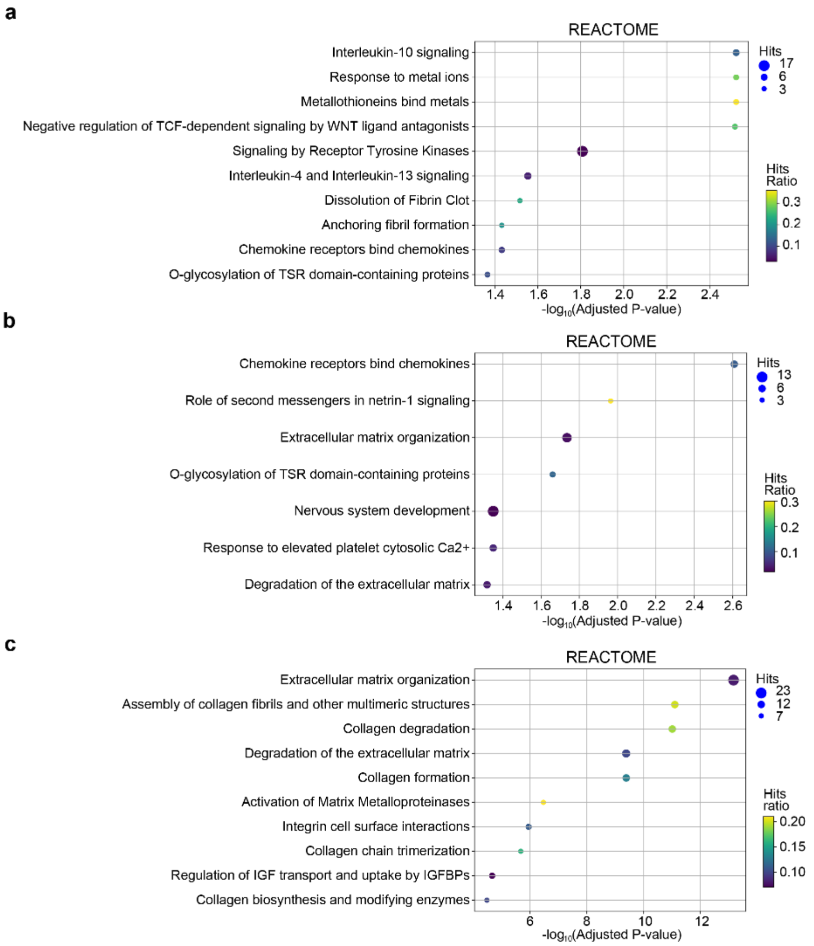


**Supplementary Fig. 11** REACTOME enrichment results of the three GEPs in the lung fibroblast and myofibroblast cells. **a**, **b**, **c** Top enriched terms significantly upregulated genes in GEP 1 (**a**), GEP 2 (**b**) and GEP 3 (**c**), respectively.

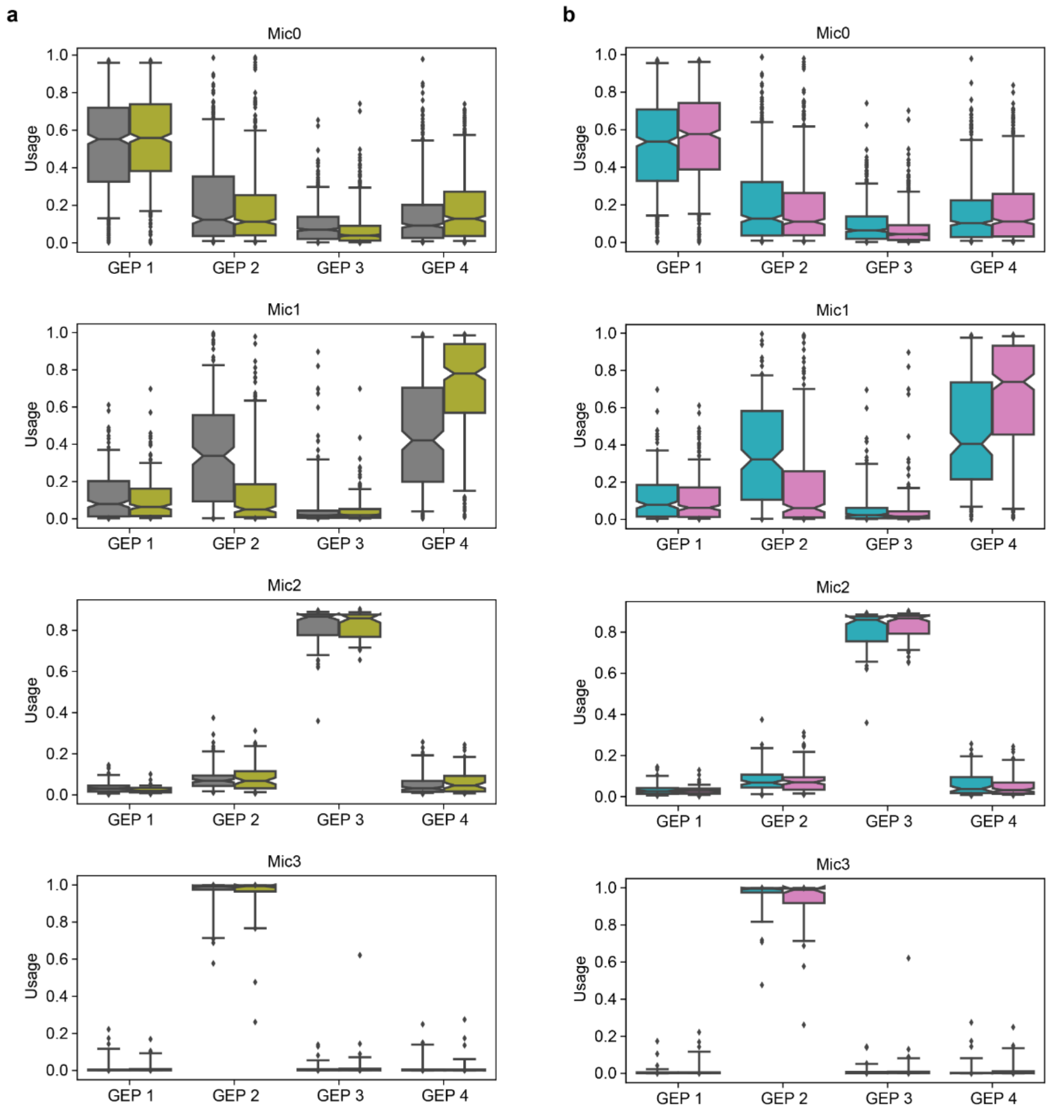


**Supplementary Fig. 12** **a**, **b** Box and whisker plot of the usage of each GEP in cells of subcluster Mic0, Mic1, Mic2, and Mic3 (from top to bottom), colored by AD pathology group and by sex in **a** and **b**, respectively. AD group and sex are color-coded in the same way as in Fig. 9b and c. Central lines represent medians, boxes represent the IQR, and whiskers represent the 5th and 95th quantiles.

**Supplementary Tables**

**Supplementary Table 1.** Top genes identified in each GEP in the pancreatic islet dataset

**Supplementary Table 2.** Upregulated DEGs (adjusted p-value < 0.05) of GEP 2 in the pancreatic islet dataset

**Supplementary Table 3.** Upregulated DEGs (adjusted p-value < 0.05) of GEP 7 in the pancreatic islet dataset

**Supplementary Table 4.** Enriched GO biological process terms of GEP 2 in the pancreatic islet dataset

**Supplementary Table 5.** Enriched Reactome terms of GEP 2 in the pancreatic islet dataset

**Supplementary Table 6.** Enriched GO biological process terms of GEP 7 in the pancreatic islet dataset

**Supplementary Table 7.** Enriched Reactome terms of GEP 7 in the pancreatic islet dataset

**Supplementary Table 8.** Upregulated DEGs (adjusted p-value < 0.05) of GEP 1 identified only using beta cells in the pancreatic islet dataset

**Supplementary Table 9.** Upregulated DEGs (adjusted p-value < 0.05) of GEP 2 identified only using beta cells in the pancreatic islet dataset

**Supplementary Table 10.** Top genes identified in each GEP in the lung fibroblast and myofibroblast cells

**Supplementary Table 11.** Upregulated DEGs (adjusted p-value < 0.05) of GEP 1 in the lung fibroblast and myofibroblast cells

**Supplementary Table 12.** Upregulated DEGs (adjusted p-value < 0.05) of GEP 2 in the lung fibroblast and myofibroblast cells

**Supplementary Table 13.** Upregulated DEGs (adjusted p-value < 0.05) of GEP 3 in the lung fibroblast and myofibroblast cells

**Supplementary Table 14.** Enriched GO biological process terms of GEP 1 in the lung fibroblast and myofibroblast cells

**Supplementary Table 15.** Enriched Reactome terms of GEP 1 in the lung fibroblast and myofibroblast cells

**Supplementary Table 16.** Enriched GO biological process terms of GEP 2 in the lung fibroblast and myofibroblast cells

**Supplementary Table 17.** Enriched Reactome terms of GEP 2 in the lung fibroblast and myofibroblast cells

**Supplementary Table 18.** Enriched GO biological process terms of GEP 3 in the lung fibroblast and myofibroblast cells

**Supplementary Table 19.** Enriched Reactome terms of GEP 3 in the lung fibroblast and myofibroblast cells

**Supplementary Table 20.** Upregulated DEGs (adjusted p-value < 0.05) of GEP 1 in the microglia of the prefrontal cortex dataset

**Supplementary Table 21.** Upregulated DEGs (adjusted p-value < 0.05) of GEP 2 in the microglia of the prefrontal cortex dataset

**Supplementary Table 22.** Upregulated DEGs (adjusted p-value < 0.05) of GEP 3 in the microglia of the prefrontal cortex dataset

**Supplementary Table 23.** Upregulated DEGs (adjusted p-value < 0.05) of GEP 4 in the microglia of the prefrontal cortex dataset

**Supplementary Table 24.** Enrichment of DEGs in each GEP across sets of makers of four microglia subclusters

**Supplementary Table 25.** Enriched GO biological process terms of GEP 2 in the microglia of the prefrontal cortex dataset

**Supplementary Table 26.** Enriched GO biological process terms of GEP 3 in the microglia of the prefrontal cortex dataset

**Supplementary Table 27.** Enriched GO biological process terms of GEP 4 in the microglia of the prefrontal cortex dataset

**Supplementary Notes**

### 1. Archetypal Analysis

Linear archetypal analysis has its mathematical definition that was first brought out by Adele Cutler and Leo Breiman in 1994[4]. For multidimensional data $\boldsymbol{x}_{i=1,\dots,n}$, where each $\boldsymbol{x}_i$ is a vector of length p, the goal of archetypal analysis is to find K archetype vectors $\boldsymbol{z}_{k=1,\dots,K}$ such that the data can be well approximated by convex combinations of the K archetypes. This is equivalent to minimizing the following loss function

$$\sum_{i=1}^{n} \|\boldsymbol{x}_i - \sum_{k=1}^{K} a_{ik}\boldsymbol{z}_k\|, \text{ where } a_{ik} \geq 0 \text{ and } \sum_{k=1}^{K} a_{ik} = 1. \quad \text{(S.1)}$$

However, minimizing equation (S.1) alone does not guarantee that the solution for $\boldsymbol{z}_{k=1,\dots,K}$ is unique, because we can find coefficient vectors $\boldsymbol{a}_{i=1,\dots,n}$ making equation (S.1) equal zero as long as $\boldsymbol{z}_{k=1,\dots,K}$ locate outside the convex hull of the data. Based on the purpose of archetypal analysis, archetypes represent 'pure' types in the data and they should resemble the real-world data. Consequently, Adele Cutler and Leo Breiman proposed that archetypes fall on the convex hull of the data so that they are extreme data values. Mathematically, this means

$$\boldsymbol{z}_k = \sum_{i=1}^{n} b_{ki}\boldsymbol{x}_i, \text{ where } b_{ki} \geq 0, \sum_{i=1}^{n} b_{ki} = 1 \text{ and } k = 1, \dots, K. \quad \text{(S.2)}$$

Note that $\boldsymbol{b}_{k=1,\dots K}$ is another set of coefficient vectors for archetypes in terms of data values. Minimizers of equation (S.1) satisfying equation (S.2) are archetypes of the data. These two conditions go hand in hand with each other to make sure that the data can be approximated by mixtures of archetypes that are convex combinations of the data values $\boldsymbol{x}_{i=1,\dots,n}$.

As we discussed in the main text, linear archetypal analysis is not suitable for systems where data values are generated non-linearly from the combination of archetypes, such as scRNA-seq data we focus. A new framework for non-linear archetypal analysis has been proposed based on autoencoders, which has been widely used for non-linear decomposition. The main challenge we face when using autoencoders for archetypal analysis is to consider how to modify the structure of traditional autoencoders to satisfy the two conditions of archetypal analysis in equation (S.1) and (S.2).

The output of the encoder side, matrix $A$, is a latent representation of the data and its row vectors correspond to the coefficient vectors $\boldsymbol{a}_{i=1,\dots,n}$ in equation (S.1). Therefore, matrix $A$ should be non-negative and have a constant row sum as 1. This constraint can be easily achieved by applying Softmax transformation on the output of the encoder. However, equation (S.2) is harder to satisfy because the non-linear transformation makes the locations of archetypes in the latent space relatively arbitrary. If we take a close look at condition (S.2), we will find what it does is to make inferred archetypes tight to the data. Therefore, one solution is to set the positions of archetypes in the latent space as a priori and penalize the distance between the preset archetypes and inferred archetypes in the latent space. To realize this, we make the encoder not only output the coefficient matrix $A$ but also another coefficient matrix $B^T$ whose column vectors correspond to $\boldsymbol{b}_{k=1,\dots,K}$ in (S.2). Then, we can add an archetypal constraint as a new loss term in equation (3) shown in the main text.
